# A scoping review of health-based survey instruments validated in Brunei Darussalam

**DOI:** 10.1101/383547

**Authors:** Mohammed M. Alhaji, Jackson Tan, Lin Naing, Nik AA Tuah

## Abstract

This study sought to map and review validated health-based survey instruments in Brunei Darussalam. A scoping search of relevant articles was carried out. Six health-based survey tools have been psychometrically evaluated in Brunei Darussalam, 4 in Brunei-Malay (SF-36v2, EQ-5D/VAS, CPQπ__14_, and m-SEQ-12) and 2 in English (OFER and WPBA) languages. Two studies (m-SEQ-12, CPQ_11–14_) translated tools in English into Brunei-Malay. Two studies (SF-36v2, EQ-5D) cross-culturally adapted the Malaysian and Singaporean versions of the tools into Brunei-Malay. Four studies were adult- and hospital-based, among healthcare workers (OFER,WPBA) and patients with chronic diseases (SF-36v2, EQ-5D); and 2 studies (m-SEQ-12, CPQ_11–14_) were non-adult-and secondary school-based. Pretesting was carried out in 4 studies (SF-36v2, EQ-5D, CPQ_11–14_, and m-SEQ-12) on a sample of 5 to 20 volunteers. The sample size for validation ranged from 40 to 457. Reliability tests, Cronbach’s alpha and intra-class coefficient (n=3), Cohen’s Kappa (n=1), and 5-point scale qualitative assessment (n=1) were measured. Validity tests included face validity (n=2), discriminant validity (n=2), convergent validity (n=2), construct validity (n=2), factorial validity (n=2), and 5-point scale qualitative assessment (n=1). There is a need for more psychometric evaluation of questionnaires in Brunei Darussalam. Importantly, large heterogeneous participants, more languages, and varied psychometric tests should be considered.

## Introduction

The use of standardized and validated questionnaires in health research and clinical practice has gained great prominence over the years [1]. Disease-specific and generic questionnaires are increasingly being used in clinical trials, disease management, and population health monitoring [2]. However, the majority of the health questionnaires have been developed in the English language[3], thus, their applicability in culturally different population or language requires cross-cultural adaptation for lexical, contextual equivalence, and psychometric tests for reliability and validity [4].

The tools translated and adapted for Asian population have shown high psychometric variation from their original versions developed in western population, due to cultural and language peculiarities [5]. Several health-based questionnaires have been translated into the Malaysian Malay which has up to 20% lexical difference from the Brunei-Malay [6]. However, health researchers in Brunei Darussalam often overlook the important lexical distinction between the two dialects and the potential psychometric dimensional variances that remain unadjusted in adopting Malaysian-Malay health, or self-developed, survey tools with little or no psychometric evaluation [7].

Thus, owing to the ever-increasing application of the scoping review method [8], this review was carried out to map the psychometrically validated health-based survey tools in Brunei Darussalam, and make recommendations for future questionnaire-based studies.

## Methods

### Search Strategy

A broad and systematic search of the literature, using medical subject heading (MeSH) and other relevant search terms, was carried out across PUBMED, MEDLINE and EBSCOhost databases to identify relevant published articles. A search in other databases such as Scopus and Google Scholar, and hand search for relevant grey literature, was also carried out. The search was limited to articles published in English language and no date was specified.

### Study Inclusion Criteria

A study was included if it met all the inclusion criteria: (a) the study is original, (b) it psychometrically evaluated an adapted, or a locally-developed, health-based questionnaire in the Bruneian population, (c) and was published in the English language. The studies which psychometrically evaluated questionnaires deemed non-health-based in the Bruneian population were excluded. An health-based questionnaire, in this review, was defined as any tool which measured any concept of health or disease, as defined by the World Health Organization [9].

### Studies and data extraction

One author searched for the articles in the databases, another author scrutinized the retrieved articles for inclusion criteria. The variables of interest were extracted using a modified Systematic Scoping Review protocol, adapted from the standard Joanna Briggs Institute (JBI)Systematic reviews protocol [8]. Overall, information on psychometric evaluation (cultural adaptation, validity and reliability tests), the type of questionnaire, questionnaire item, sample size, sample demographics, and study conclusion were retrieved. The retrieved articles were narratively reviewed, and no statistical procedure was carried out.

## Results

### Study Search and Selection

A total of 401 articles were returned from the database. After the screening of the returned articles, 14 were retained. A further scrutiny of the selected articles against the inclusion criteria resulted in the exclusion of 8 articles. In the end, 6 studies were included in this review (Fig. 1).

**Figure 1:**
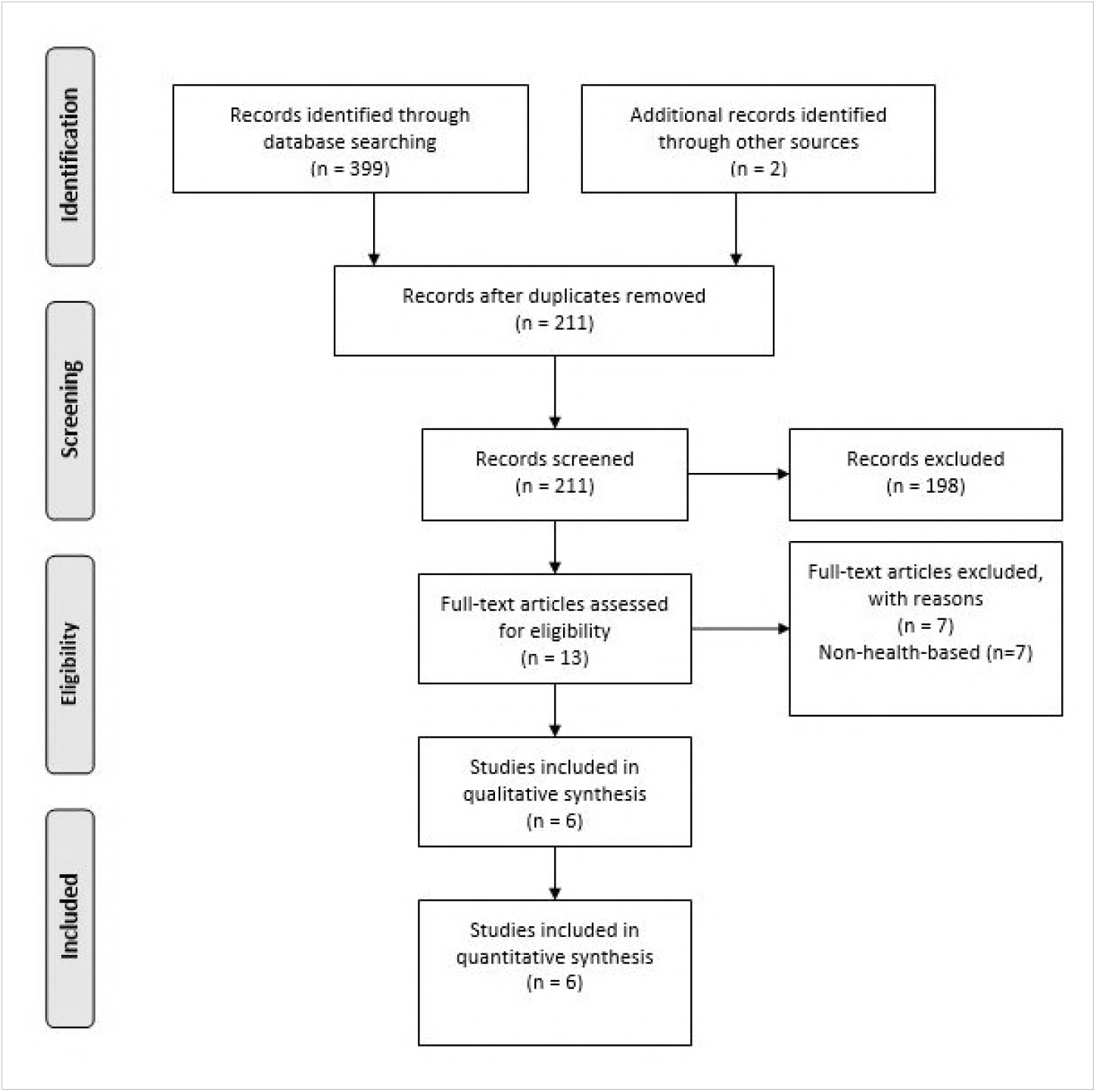
PRISMA Study Search and Selection Flow Diagram[10]

### Characteristics of the studies

The studies (n=6) were all cross-sectional in design, with non-probabilistic sampling—convenient (n=3), purposive (n=3)—being the commonest sampling method. One study used random and convenient sampling methods.[11] Sample size ranged from 40 [12] to 457 [13]. Four studies—Short Form version 2, SF-36v2 [11], EuroQol/Visual Analogue Scale, EQ-5D/VAS [6], Child Perception Questionnaire, CPQ_11-14_ [13], and modified-Smoking Self-Efficiency Questionnaire, m-SEQ-12 [12]—evaluated tools adapted into Brunei-Malay while 2 studies— Occupational Fatigue Exhaustion Recovery, OFER [14], and Workplace-Based Assessment, WPBA [15]— validated English-based tools. One study was a feedback on postgraduate medical training assessments (WPBA) [15].

Four studies [6, 11, 14, 15] were conducted on adults (18 years old and above), and two studies [12, 13] were conducted on children and adolescents (less than 18 years old). Two studies [12, 13] were school-based (among high school students), and four studies were hospital-based, among healthcare workers (n=2) [14, 15] and patients with chronic disease (n=2) [6, 11]. One studies had a heterogeneous sample, individuals from the general population and patients with chronic kidney disease [11]. (Table 1)

### Psychometric procedure and tests

Two studies translated the tools in the English language to Brunei-Malay using forward and backward translation method [12, 13]. Two studies cross-culturally adapted the tools for Bruneian context from the neighbouring Malaysian Malay [6, 11] and Singaporean Malay [6]. Tool pretesting or piloting on a sample size ranging from 5 to 20 was carried out in four studies[6, 11, 12, 13].

On reliability test, internal consistency or homogeneity (Cronbach’s alpha) were used in 3 studies [11, 12, 13]; test-retest or stability (intra-class coefficient) in 3 studies [6, 12, 13], Kohen’s Kappa in one study [6], and 5-point scale qualitative feedback in one study [15]. A wide variety of validity tests were used. Face validity (n=2) [11, 14], discriminant validity (n=2) [11, 13], convergent validity (n=2) [11, 14], construct validity (n=2) [12, 13], factorial validity (n=2) [11, 12], and 5-point scale qualitative feedback (n=1) [6] were used in determining validity of the tools (Table 1).

**Table 1.**
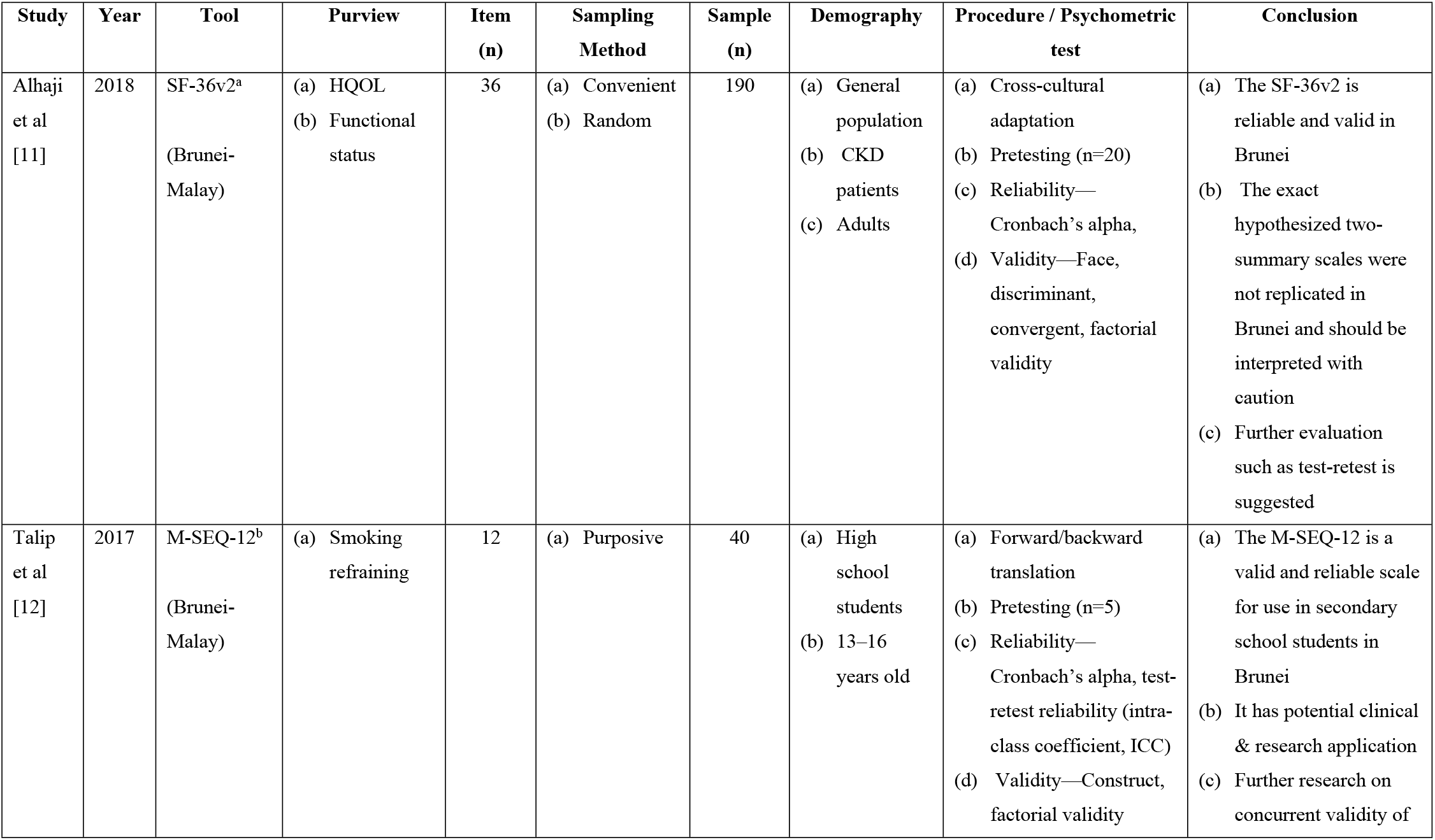
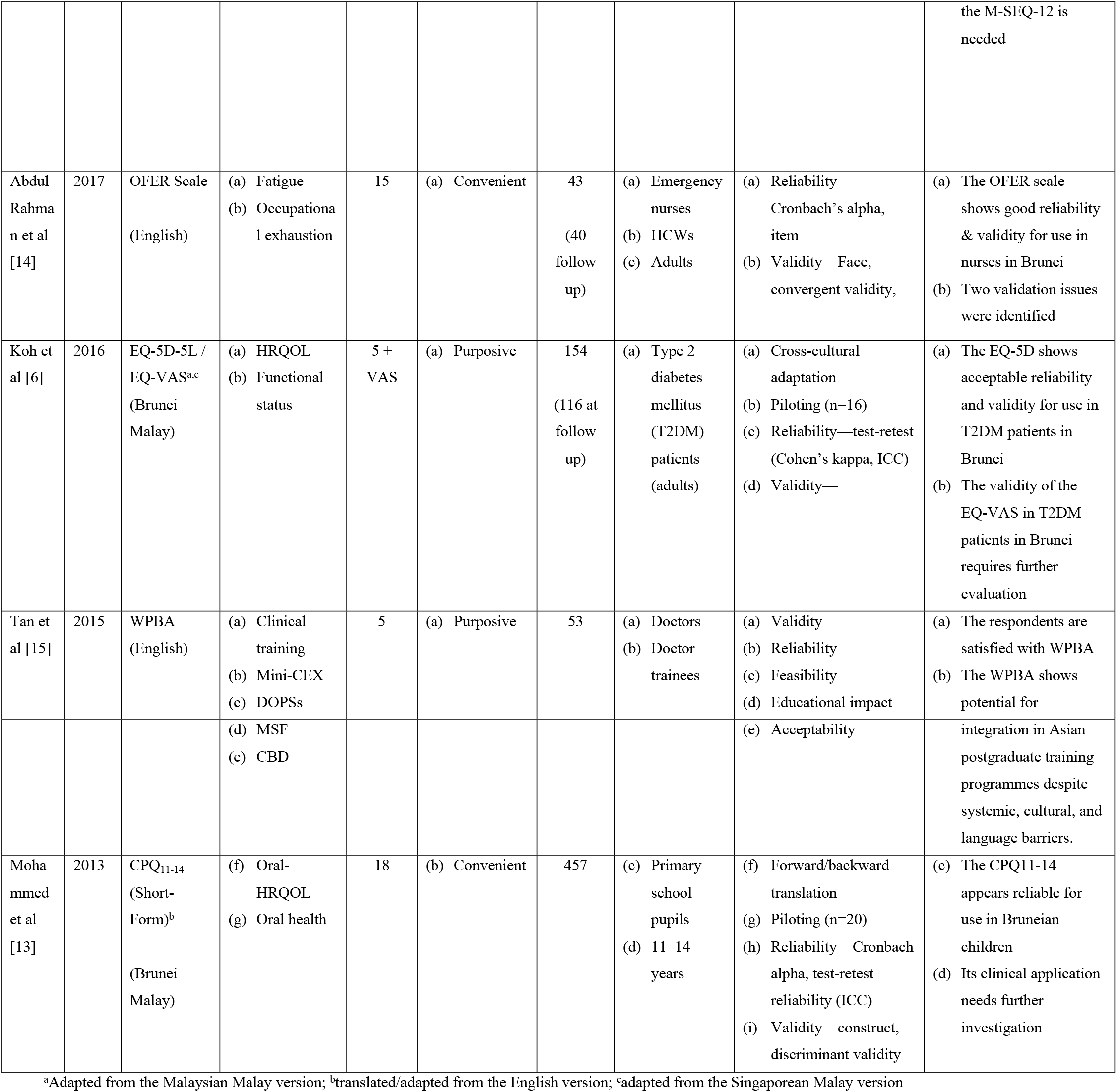
Description of Health-based health survey validated in Brunei Darussalam.

| Study | Year | Tool | Purview | Item (n) | Sampling Method | Sample (n) | Demography | Procedure / Psychometric test | Conclusion |
| --- | --- | --- | --- | --- | --- | --- | --- | --- | --- |
| Alhaji et al [11] | 2018 | SF-36v2<sup>a</sup><br>(Brunei-Malay) | (a) HQOL<br>(b) Functional status | 36 | (a) Convenient<br>(b) Random | 190 | (a) General population<br>(b) CKD patients<br>(c) Adults | (a) Cross-cultural adaptation<br>(b) Pretesting (n=20)<br>(c) Reliability—Cronbach's alpha,<br>(d) Validity—Face, discriminant, convergent, factorial validity | (a) The SF-36v2 is reliable and valid in Brunei<br>(b) The exact hypothesized two-summary scales were not replicated in Brunei and should be interpreted with caution<br>(c) Further evaluation such as test-retest is suggested |
| Talip et al [12] | 2017 | M-SEQ-12<sup>b</sup><br>(Brunei-Malay) | (a) Smoking refraining | 12 | (a) Purposive | 40 | (a) High school students<br>(b) 13–16 years old | (a) Forward/backward translation<br>(b) Pretesting (n=5)<br>(c) Reliability—Cronbach's alpha, test-retest reliability (intra-class coefficient, ICC)<br>(d) Validity—Construct, factorial validity | (a) The M-SEQ-12 is a valid and reliable scale for use in secondary school students in Brunei<br>(b) It has potential clinical & research application<br>(c) Further research on concurrent validity of the M-SEQ-12 is needed |
| Abdul Rahman et al [14] | 2017 | OFER Scale (English) | (a) Fatigue<br>(b) Occupational exhaustion | 15 | (a) Convenient | 43<br>(40 follow up) | (a) Emergency nurses<br>(b) HCWs<br>(c) Adults | (a) Reliability—Cronbach's alpha, item<br>(b) Validity—Face, convergent validity, | (a) The OFER scale shows good reliability & validity for use in nurses in Brunei<br>(b) Two validation issues were identified |
| Koh et al [6] | 2016 | EQ-5D-5L / EQ-VAS<sup>a,c</sup> (Brunei Malay) | (a) HRQOL<br>(b) Functional status | 5 + VAS | (a) Purposive | 154<br>(116 at follow up) | (a) Type 2 diabetes mellitus (T2DM) patients (adults) | (a) Cross-cultural adaptation<br>(b) Piloting (n=16)<br>(c) Reliability—test-retest (Cohen's kappa, ICC)<br>(d) Validity— | (a) The EQ-5D shows acceptable reliability and validity for use in T2DM patients in Brunei<br>(b) The validity of the EQ-VAS in T2DM patients in Brunei requires further evaluation |
| Tan et al [15] | 2015 | WPBA (English) | (a) Clinical training<br>(b) Mini-CEX<br>(c) DOPSs | 5 | (a) Purposive | 53 | (a) Doctors<br>(b) Doctor trainees | (a) Validity<br>(b) Reliability<br>(c) Feasibility<br>(d) Educational impact | (a) The respondents are satisfied with WPBA<br>(b) The WPBA shows potential for |
|  |  |  | (d) MSF<br>(e) CBD |  |  |  |  | (e) Acceptability | integration in Asian postgraduate training programmes despite systemic, cultural, and language barriers. |
| Mohammed et al [13] | 2013 | CPQ<sub>11-14</sub> (Short-Form)<sup>b</sup><br>(Brunei Malay) | (f) Oral-HRQOL<br>(g) Oral health | 18 | (b) Convenient | 457 | (c) Primary school pupils<br>(d) 11–14 years | (f) Forward/backward translation<br>(g) Piloting (n=20)<br>(h) Reliability—Cronbach alpha, test-retest reliability (ICC)<br>(i) Validity—construct, discriminant validity | (c) The CPQ11-14 appears reliable for use in Bruneian children<br>(d) Its clinical application needs further investigation |

<sup>a</sup>Adapted from the Malaysian Malay version; <sup>b</sup>translated/adapted from the English version; <sup>c</sup>adapted from the Singaporean Malay version

HCWs, Healthcare Workers; HRQOL, Health-Related Quality of Life; SF-36, Short Form Survey-36; M-SEQ-12, Modified Smoking Self-Efficiency Questionnaire; OFER, Occupational Fatigue Exhaustion Recovery Scale; EQ-5D-5L/VAS, EuroQol/Visual Analogue Scale; WPBA, Workplace-Based Assessment; CPQ_11–14_, Child Perception Questionnaire; Mini-CEX, Mini-Clinical Examination; DOPSs, Directly Observed Practical Skills; MSF, Multisource Feedback; CBD, Case-Based Discussion

### Health-related Tools validated in Brunei Darussalam

Three studies [6, 11, 13] validated health-related quality of life (HRQOL) instruments. The SF-36v2 is a 36-item tool which measures generic HRQOL in 8 scales (physical functioning, role-physical, bodily pain, general health, vitality, social functioning, role-emotional, and mental health). These 8 domains are reducible to two summary scales, physical component summary and mental component summary. The SF-36v2 was found valid and reliable for use in Brunei, among chronic kidney disease (CKD) patients and the general population [11].

The EQ-5D or EQ-5D-5L is a 5-item utility-based tool measures generic HRQOL under 5 dimensions (anxiety/depression, mobility, pain/discomfort, self-care, and usual activities) alongside a 20cm vertical EQ-VAS with number scale between 0 and 100 [6]. The EQ-5D showed acceptable reliability and validity for use among Type 2 Diabetes Mellitus (T2DM) patients. The CPQ_11–14_ (short form) is an 18-item instrument variant for measuring oral-HRQOL in 5 broad domains (oral health and oral health-related well-being, oral symptoms, functional limitations, emotional well-being, and social well-being) [13]. The CPQ_11-14_ was found to be reliable for use among children in Brunei.

The other 3 health-based instruments validated in Brunei bordered on smoking refraining (n=1),[12] occupational exhaustion and fatigue among healthcare workers (HCWs) (n=1) [14], and workplace-based assessment of doctor trainees [15]. The m-SEQ-12 is a 12-item scale which measures one’s capability to desist from smoking in two dimensions (internal and external stimuli) [12]. The m-SEQ-12 was found to be valid and reliable for use among secondary school students and have potential clinical and research application in Brunei Darussalam.

The OFER scale is a 15-item scale for measuring work-related fatigue under 3 domains (chronic fatigue, acute fatigue, and persistent fatigue) [14]. The OFER scale showed good reliability and validity for use among nurses in Brunei. The WPBA tools (Mini-Clinical Examination, Directly Observed Practical Skills, Multisource Feedback, and Case-Based Discussion) were evaluated among doctors and doctor trainees for “validity, reliability, feasibility, educational impact, and acceptability” [15]. The findings showed excellent satisfaction in all the 5 headings, thereby showing potential for integration in postgraduate medical training in Asia.

However, further investigations were suggested in the studies. Additional psychometric evaluation such as the test-retest validity test and caution in interpreting the two-dimensional summary scores of the SF-36v2 have been advised [11]. Other recommendations include concurrent validity test for m-SEQ-12 [12]; increased sample size and convergent validity test, by inference, for the OFER scale [14]; further validity tests on EQ-VAS in T2DM patients [6]; further investigation on clinical application of the CPQ_11–14_ [13]; and systemic, cultural and language barrier concerns for the WPBA [15].(Table 1)

## Discussions

A lot of questionnaire-based studies in Brunei Darussalam either adapt existing questionnaires, often from Malaysia or used directly translated tools or self-developed questionnaires with little or no psychometric evaluation [7]. The use of non-or poorly-validated questionnaires affect data quality, outcomes and comparability of the finding with other similar findings.[2, 16]. Our review showed that Cronbach’s alpha test was the most widely used test of reliability of quantitative tool validation in Brunei Darussalam, which is similar to what is obtainable globally [17]. However, its ubiquitous use has been criticized for being misunderstood and misapplied [18], thereby leading to the suggestion that Cronbach’s alpha should be measured alongside other tests of reliability [19] such as the split-half [20].

The majority of the tools (n=4) were validated in the Malay language and none in Chinese or other minority languages. Ethnic-wise, Brunei Darussalam is composed of Brunei Malay Brunei Malay (65.7%), Chinese (10.3%), and others (24.0%) which comprised of the indigenous groups of the Malay race namely Belait, Bisaya, Brunei, Dusun, Kedayan, Murut and Tutong [21]. Although Malay is the lingua franca in Brunei Darussalam, the validations of survey tools in the other minority languages, especially Chinese language, are need needed, in line with the international best practice of survey tools cross-cultural adaptation [22].

Similarly, the majority of the health-based tools evaluated in Brunei Darussalam were validated on a homogenous sample rather than a heterogeneous sample, thereby making such tool inapplicable for certain population which were not covered in the validation [16]. For a tool to be applicable in a broad population, its whole psychometric scale length needs to be validated in a heterogeneous sample from a wide range of sociodemographic strata (e.g. a mix of the old and the young, sick and healthy, male and female) rather than in a homogeneous sample, as the latter lower variance and factor loading [23]. Further, the sample sizes for the psychometric evaluations of health-based tools in Brunei Darussalam were usually small. The minimum sample size recommended for factor analysis, an important validity test, was 300 participants [23, 24]. The highest sample size for health-based validation in Brunei Darussalam was 457 participants [13].

This was often not the case for other psychometric evaluations of non-health-based tools validated in Brunei Darussalam. Example, the My Class Inventory (MCI) tool for assessing ‘students’ perceptions of the classroom learning environment (n=1565) [25], the What Is Happening In This Class (WIHIC) for measuring ‘perceptions of learning environment’ (n=644) [26], the Students’ Perception of Assessment Questionnaire (SPAQ) for measuring students perception of assessment (n=1028) among secondary school students [27], the Questionnaire on Teacher Interaction (QTI) for evaluating ‘students’ Enjoyment of their Sciences Lessons (ENJ)’ among students among primary school pupils (n=3104) [28], and the Cultural Learning Environment Questionnaire (CLEQ) for evaluating ‘culturally-sensitive’ factors in teacher trainees learning environments’ among trainee teachers (n=475) [29].

Three other non-health validations in Brunei Darussalam with low sample size included the Eysenck Personality Questionnaire-Revised (EPQ-R) Short Scale for measuring personality traits among trainee teachers (n=223) [30], the ‘MODEL (Meanings, Organise, Develop, Execute, Link)’, a local framework, for ‘exploring students’ cognitive competency in solving non-routine problems’ among junior college students (n=167) [31], and the Minnesota Multiphasic Personality Inventory-Revised (MMPI-2) for assessing personality traits among student teachers (n=84) [7]. Although fewer than 100 participants have been used to psychometrically evaluate a tool,[32] a sample size of 100 and below is considered ‘poor sampling’[24]. The use of convenience sampling method has also been considered a limitation for generalizability [16].

Finally, our scoping review of health-based instruments validated in Brunei Darussalam highlights the paucity of published psychometric evaluation of health-based tools, the use of small and homogenous sample sizes, the lack of diversity in the language of the tools, and the absence of diversity of psychometric tests used in the validation processes. We, therefore, recommend that health and medical researchers in Brunei Darussalam to endeavour to validate and publish the questionnaires they adopt or self-develop before use in research. Importantly, the psychometrically evaluation of the tools should aim to include large heterogeneous participants and the languages should be diversified to include other languages such as the Chinese language. Similarly, varied sampling strategies and psychometric tests should be used for survey tool validations.

